# GM1a functions as a coreceptor/ attachment factor for Dengue virus during infection in mammalian systems

**DOI:** 10.1101/2022.01.20.477180

**Authors:** Sarala Neomi Tantirimudalige, Palur Venkata Raghuvamsi, Jonathan Chua Wei Bao, Ganesh S. Anand, Thorsten Wohland

## Abstract

Dengue virus (DENV) is a flavivirus causing an estimated 390 million infections per year around the world. Despite the immense global health and economic impact of this virus, its true receptor(s) for internalization into live cells has not yet been identified, and no successful antivirals or treatments have been isolated to this date. This study aims to improve our understanding of virus entry routs by exploring the sialic acid-based cell surface molecule GM1a and its role in DENV infection. The interaction of the virus with GM1a was studied using fluorescence correlation spectroscopy (FCS), fluorescence cross correlation spectroscopy (FCCS), imaging FCS (ImFCS) and amide hydrogen/deuterium exchange mass spectrometry (HDXMS), and the effect on infectivity and movement of the virus during infection was explored using plaque assays and fluorescence-based imaging and single particle tracking (SPT). GM1a was deemed to interact with DENV at domain I (DI) and domain II (DII) of the E protein of the protein coat at quaternary contacts of a fully assembled virus, leading to a ten-fold increase and seven-fold increase in infectivity for DENV1 and DENV2 in mammalian cell systems respectively. The interaction of virus with GM1a triggers a speeding up of virus movement on live cell surfaces, possibly resulting from a reduction in rigidity of cellular rafts during infection, and functions as a coreceptor/ attachment factor for DENV during infection in mammalian systems.

**Author Summary:** Dengue virus (DENV) is a flavivirus causing an estimated 390 million infections per year around the world. Despite the immense global health and economic impact of this virus, no successful antivirals or treatments have been isolated to this date. This may be due to the incomplete understanding of the virus infection mechanism, including a lack of an identified ‘true’ receptor and entry related attachment factors or co-receptors responsible for internalization of the virus. This work focuses on the early infection stage of DENV1 and DENV2 strains, to identify how the virus moves on cell surfaces in its search for its receptors, and identifies the critical role of the sialic acid ganglioside GM1a during internalization of the virus.

## Introduction

Dengue virus (DENV) is a mosquito-borne enveloped virus, from the *Flaviviridae* family. It is transmitted from human to human by the bite of an infected mosquito (*Aedes aegypti*, and occasionally *Ae. albopictus*) with symptoms ranging from fever, muscle and joint pain (Dengue fever) to a more life-threatening haemorrhagic fever or shock syndrome, in both adults and children alike [1–3]. It has been estimated that there are as many as 390 million dengue infections per year, spread throughout 128 countries, of which 96 million manifest clinically with symptoms [4–6], posing a major global health impact.

DENV has four identified serotypes (DENV1, DENV2, DENV3, DENV4). It has a diameter close to 500 Å with an outermost protein shell which is embedded in the host-derived lipid bilayer, which in turn encapsulates the ∼11 kb single stranded positive sense RNA genome which encodes 10 viral proteins, out of which 3 are structural (capsid (C), the pre-membrane or membrane (prM or M respectively) and the envelope (E) proteins) and 7 are non-structural (NS1, NS2a, NS2b, NS3, NS4a, NS4b and NS5) in function [1,4,7–9]. The outer most protein shell of mature DENV is made up of 180 copies of E and 180 copies of M proteins arranged in an icosahedral manner [4]. The E protein is the major antigenic structure on the surface of the virus and is involved in receptor and attachment factor binding [4,8,10–16]. Infectious DENV particles interact with attachment factors on the cell membrane, followed by movement along the cell surface in its search for its receptors and co-receptors, which internalize it into its host cell [13–15,17,18].

Current research work and literature highlight a number of receptors/co-receptors/attachment factors involved in DENV internalization, with no definitive answer on what factors are truly the most critical for virus internalization. The adhesion molecule of dendritic cells DC-SIGN, ER chaperonin GRP-78, the 37/67 kDa high-affinity laminin receptor, heat shock proteins 70 and 90, glycosaminoglycans (GAGs) such as heparan sulfate (HS), heparan sulfate proteoglycans (HSPG) and lectins, TIM and TAM proteins, mannose receptor (MR) of macrophages, lipopolysaccharide (LPS) receptor CD14, glycosphingolipids (GSL), claudin-1, are a few among these explored receptors and attachment factors for DENV in mammalian cell systems [1,12,26–28,14,19–25]. Having a broad range of receptors and entry routes, DENV appears to possess the ability to infect many different types of cells, using many different mechanisms, and may use entry routes more ubiquitous in nature and reflective of the diversity in cell surface composition.

In this work we focus on the ganglioside GM1a and describe its involvement in DENV infection in mammalian cell systems. GM1a is a glycosphingolipid, with a glycan part and lipid portion which contributes to the glycocalyx and lipidome of cells, respectively. It possesses a terminal sialic acid moiety, which interacts with cargo such as Cholera Toxin B (CTxB), viruses and bacteria, in order to internalize them into cells [29–34]. Sialic acids of the ganglioside family, are ubiquitously found in most mammalian cells and are important for cell signalling, cell adhesion and many other cellular functions [35–41]. Mammalian cells have a dense glycocalyx, and more often than not, the first point of contact for any pathogen will be the glycan interaction points on cell surfaces, and more than half of all known mammalian viruses are reported to interact with glycans during internalization. The diversity of viruses that can interact with sialic acid moieties range from membranous to non-membranous, those that have capsids, to those that do not, and both virus types with RNA encoded or DNA encoded genomes [42–48].

One such virus is the Influenza A virus (IAV) (*Orthomyxoviridae*), which shows interaction with sialic acid moieties on host cell surface during infection [44,46,48–51]. The virus surface is decorated with two surface glycoproteins, hemagglutinin (H/HA) and neuraminidase (N/NA). The hemagglutinin is responsible for interacting with the sialic acid moiety (Neu5Ac) on cell surfaces, and initiates viral internalization, possibly with the help of another more proteinaceous receptor [52–55]. Paramyxovirus (*Paramyxoviridae*) binds sialic acid residues in a similar fashion to IAV, where the virus surface hemagglutinin-neuraminidase (HN) glycoprotein binds to sialic acid on host cells, mediating virus internalization [56]. Newcastle Disease (NDV), Sendai, mumps, and parainfluenza viruses are reported to recognize glycans with the terminal NeuAca2-3Gal linkage [46, 57]. Simian virus 40 (SV40) is reported to be highly specific for GM1 binding, where the capsid protein, VP1, forms a complex with the carbohydrate portion of GM1, confirmed by crystallography studies [46,51,58–60]. Coronavirus (*Coronaviridae*) surfaces hold two glycoproteins, a spike protein and hemagglutinin-esterase, which can both interact with glycans as primary/co-receptors or as attachment factors. Feline coronavirus (FCoV), Transmissible gastroenteritis virus (TGEV), Porcine Epidemic Diarrhoea (PED), have all been reported to bind sialic acid host at cell receptors (NeuAc/NeuGc) via the spike protein as a secondary receptor [61]. The spike on Human Respiratory Coronavirus (OC43) and Human Coronavirus (HKU1), is reported to interact with NeuAc as a primary receptor, while Middle East respiratory syndrome (MERS) binds sialic acid with a preference for an α2,3-link [46,51,62–64]. Picornaviruseses (*Picornaviridae*) Coxsackie A24, human enterovirus 68 and Murine encephalomyocarditis virus, all bind sialic acid receptors at the NeuAca2-6Gal and/or NeuAca2-3Gal sites on N-glycans [46,65–67]. It has been recently shown that Sialic acid-containing glycolipids including GM1 mediates binding and viral entry of SARS-CoV-2 [68]. Various viruses in the families of Polyomavirus (*Polyomaviridae*), Parvovirus (*Parvoviridae*), Rotavirus and Orthoreovirus (*Reoviridae*), Caliciviruses (*Caliciviridae*), and Mammarena viruses in the family of Arenaviruses (*Arenaviridae*), have all been shown to interact with sialic acid containing glycans during cellular infection [44,46,69–74], making sialic acid containing glycan receptors a widely utilized entry route by many types of viruses.

In this work, the interaction of DENV1 (PVP 159) and DENV2 (NGC strain) with the sialic acid ganglioside GM1a was explored and confirmed by Fluorescence Cross Correlation Spectroscopy (FCCS), Fluorescence colocalization studies, Fluorescence Single Particle Tracking (Fluorescence SPT), and Imaging Fluorescence Correlation Spectroscopy (ImFCS). The interaction site of GM1a with DENV was mapped by amide hydrogen/deuterium exchange mass spectrometry (HDXMS). The impact on infectivity of this interaction of DENV1 and DENV2 with GM1a was tested by plaque assays, and it is shown that both DENV1 and DENV2 show increased infectivity in the presence of GM1a. Taken together our results show that, DENV1 and DENV2, interact with GM1a on mammalian cells, resulting in increased infectivity of the virus, and functions as an attachment factor/receptor/co-receptor during virus internalization.

## Results

### Interaction of Dengue virus with surface GM1a on live mammalian cells

The involvement of GM1a during DENV infection was probed using colocalization studies and quasi pulsed interleaved fluorescence cross-correlation spectroscopy (qPIE-FCCS) [75] in live Vero cells. Colocalization studies and time-lapse imaging carried out on a confocal microscope showed that DENV1 and DENV2 particles each colocalized and moved together with GM1a-Bodipy on live cell surfaces (S2 Fig and S1 Table). For a more direct measurement of interactions, qPIE-FCCS experiments on the confocal microscope were conducted between GM1a labelled with Bodipy FL (GM1a-Bodipy), and DENV labelled with Alexa Fluor 555 NHS (Fig 1). The degree of interaction in qPIE-FCCS was semi-quantitatively evaluated using the so-called *q*-value, which is proportional to the interaction detected (see materials and methods section). As negative control we used DiIC18(3) and GM1a-Bodipy, two lipids not expected to interact as they partition into liquid disordered or liquid ordered regions, respectively. As positive control we used PMT-mEGFP-mApple, a plasma membrane targeting sequence where both red and green fluorophores are linked to each other and thus should provide a maximum of cross-correlation achievable with the setup [76]. While qPIE-FCCS negative and positive controls showed *q*-values of 0.03±0.01 (Avg±SEM; cells = 6) and 0.45±0.01 (Avg±SEM; cells = 10), respectively, DENV1 and DENV2 showed intermediate *q*-values of 0.15±0.02 (Avg±SEM; cells = 19) and 0.21 ±0.02 (Avg±SEM; cells = 58), indicating clear interactions. As a further control we measured the interaction of GM1a and CTxB, a known GM1a binding protein [29, 30], which yielded a *q* value of 0.25±0.04 (Avg±SEM; cells = 9).

**Fig 1.**
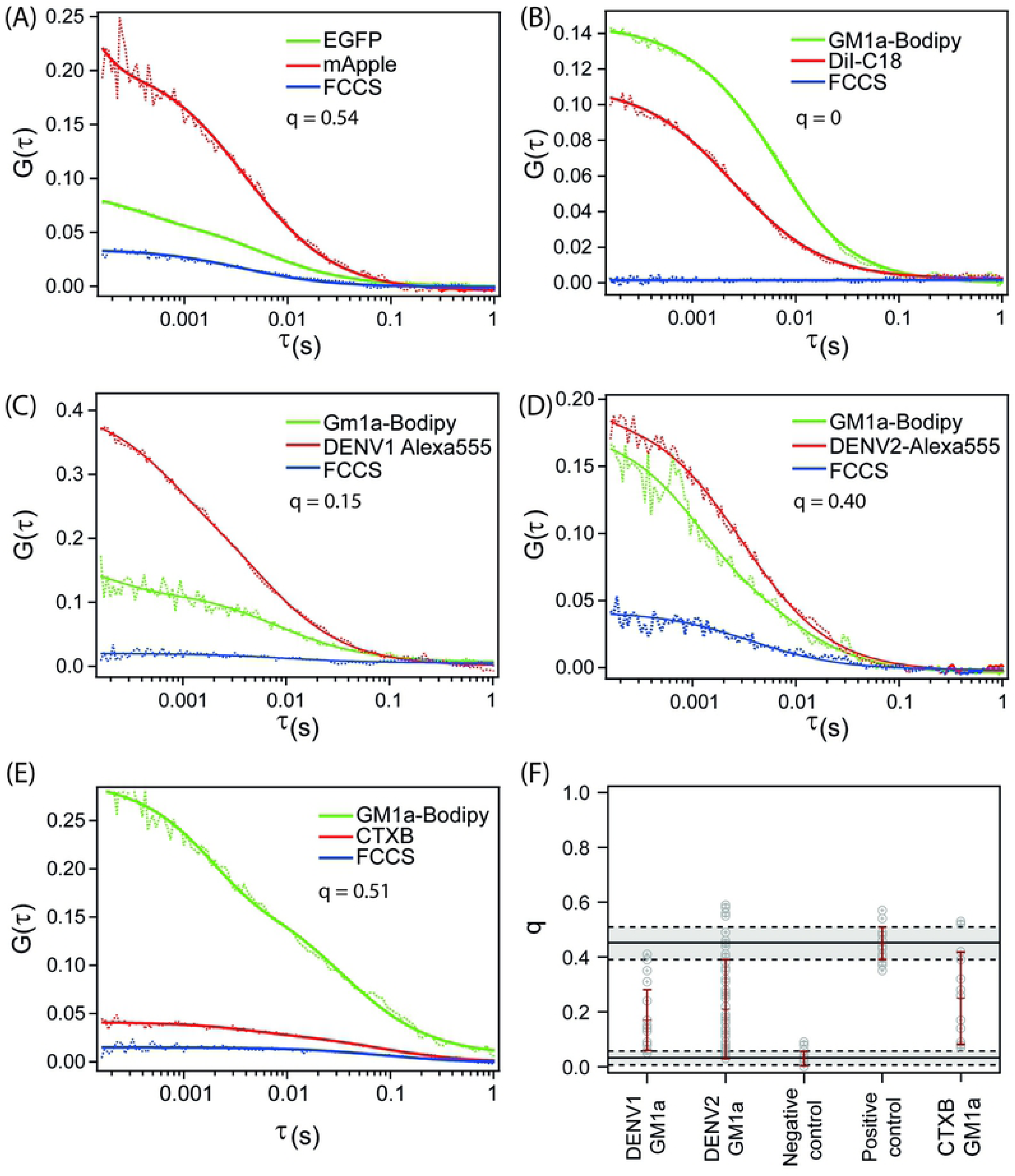
**Representative FCCS curves obtained by quasi-PIE FCCS.** (A) FCCS positive control of PMT-mEGFP-mApple with a q value of 0.54. (B) FCCS negative control of non-interacting partners GM1a-Bodipy and DiI-C18 with a q value of 0. (C) GM1a interaction with DENV1 with a q value of 0.15 (D) GM1a interaction with DENV2 with a q value of 0.40. (E) Interaction of GM1a with CTXB with a q value of 0.51 (F) q value distribution of DENV1 and DENV2 interaction with GM1a compared with the negative and positive controls. Overall q values of 0.15±0.02 (cells = 19), 0.21 ±0.02 (cells = 58), 0.45±0.01 (cells = 10), 0.03±0.01 (cells = 6) for DENV1, DENV2, positive control of PMT-mEGFP-mApple and negative control with DiIC18 (Avg±SEM). Error bars represent the standard deviation. All cells were pre-treated with DPDMP to inhibit endogenous GM1a production.

### Mapping the binding hotspots of GM1a on DENV E-protein

Colocalization of GM1a glycosphingolipid with DENV2 particles was confirmed by FCCS. We performed HDX-MS on free DENV2 and DENV2 in the presence GM1a sugar moiety to identify the binding hotspot of GM1a on the DENV viral surface. 36 pepsin proteolyzed peptides were obtained with high signal to noise ratios and covering 63% of the E protein sequence. Here, we used high molar ratio GM1a sugar moiety to that of E protein dimer (125: 1) to achieve saturation in GM1a binding on viral surface.

Comparative deuterium exchange analysis was performed between free and GM1a bound DENV2 using deuterium exchange difference plot. A deuterium exchange difference plot (Fig 2A) shows the difference in number of deuterons exchanged between DENV2:GM1a and free DENV2 at 1 min of deuterium labelling time for each pepsin digested peptide. In the presence of GM1a, peptide 40-57, 129-136 and 322-335 showed lower deuterium exchange (Fig 2A). Peptide 40-57 and 129-136 span the domain I (DI) and domain II (DII) of the E protein. Peptide 322-335 is localized at the five-fold symmetry axis of the virion (Fig 2D). Previous reported crystal structures of the E protein complexed with carbohydrate molecule β-octyl glucopyranoside (βOG pocket) (PDB: 1OKE) shows a similar binding site for sugar molecules spanning DI and DII [77]. We also performed control HDX-MS experiments to map interactions of GM1a with recombinant DENV2 E protein/soluble E protein (sE protein). We observed no changes in deuterium exchange in the presence of GM1a (S4 Fig). This indicates that GM1a requires quaternary contacts of a fully assembled virus to bind to the surface. Furthermore, DENV1 and DENV2 both show a ∼67% sequence similarity of the E protein (S5 Fig), therefore it can be expected that DENV1 may also bind GM1a in a similar fashion to that of DENV2.

**Fig 2.**
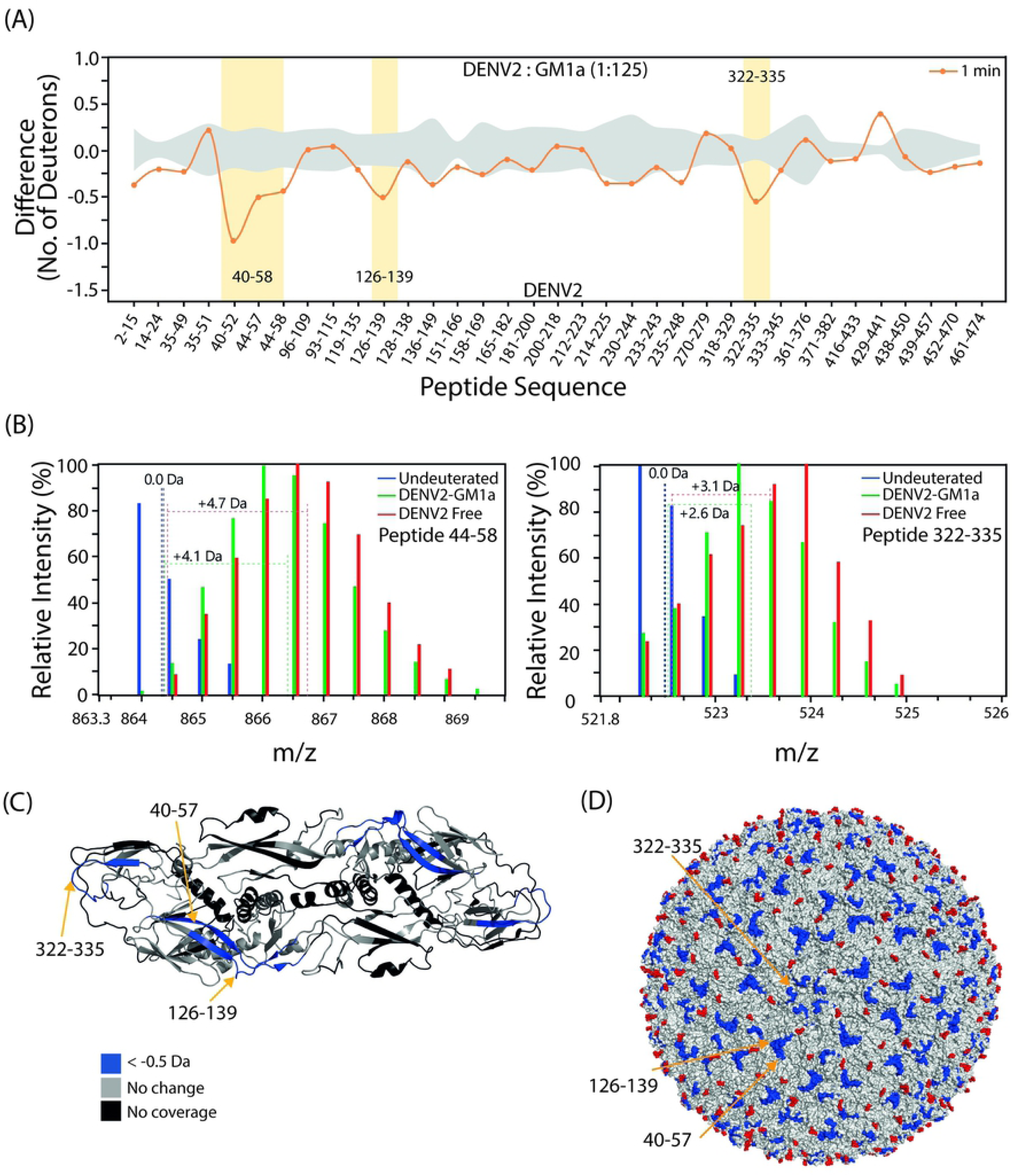
**HDX-MS reveal GM1a sugar moiety binding hotspot on Denv2 NGC surface on E protein** (A) Differences in average number of deuterons exchanged (*Y-axis*) in DENV2:GM1a state and free Denv2 state are plotted in a deuterium uptake difference plot for t=1min labelling time. Pepsin digested peptides are listed from N to C terminus (*X-axis*). Peptide regions showing high protection in the presence of GM1a are highlighted. Standard deviations are shaded gray and plots were generated using DynamX 3.0 software. The data is uncorrected for maximum deuterium content of 90% under experimental conditions, and unadjusted to accommodate loss in deuterons exchanged due to back-exchange (average back-exchange ∼15% under our HDXMS conditions). (B) Representative mass spectra of peptides protected in the presence of GM1a i.e., peptide 44-58 and 322-335 are shown and dotted lines are shown at the centroid of mass spectra (C) Differences in deuterium exchange between DENV2-GM1a bound and free DENV2 are mapped onto a E protein dimer (PDB: 3J27) with differences highlighted as per key (D) Peptides protected in the presence of GM1a sugar are mapped in blue onto the full virus (PDB:3J27). Red spheres represent glycans on viral surface.

It is important to note that the ∼37% E protein regions lacking coverage in our experiments that span domain II (peptide 59-92) and domain I/domain III (peptide 280-317) are surface exposed and form quaternary contacts in the virion. Due to the lack of deuterium exchange measurements, we cannot describe the role of these regions in binding to GM1a.

### GM1a assists and increases infection in DENV1 and DENV2

The biological significance of the proximity of GM1a and DENV observed by qPIE-FCCS and colocalization studies was further investigated on live BHK21 mammalian cells by conducting plaque assays. The infectivity levels were compared for untreated cells, GM1a depleted cells and GM1a enriched cells (Fig 3). GM1a depletion was achieved by D-PDMP treatment, while GM1a enrichment was by BSA loading (see Materials and Methods). DENV1 and DENV2 both show a similar trend in infectivity with relation to GM1a on cell surfaces, with a significant increase in infectivity seen for the GM1a enriched cells as compared to the GM1a depleted cells. DENV1 shows an increase in infectivity in GM1a enriched cells with an average value of 6.0 x10^7^ PFU/ml, as compared to 5.4 x10^6^ PFU/ml for D-PDMP treated (GM1a depleted) cells, in three repeat experiments (Fig 3A). While DENV2 shows an increase in infectivity in cells enriched with GM1a, with an average value of 1.3 x10^15^ PFU/ml versus a value of 8.5 x10^8^ PFU/ml in GM1a depleted plates, for three repeat experiments (Fig 3B). The three trials in both DENV1 and DENV2 show a similar trend of increased infectivity in GM1a enriched cells, as compared to the GM1a depleted and untreated cells. This indicates that GM1a significantly increases infectivity of DENV, but is not the only route of entry for the virus, where in the absence of GM1a, the infection is not completely abolished, and the virus internalizes by other routes.

**Fig 3.**
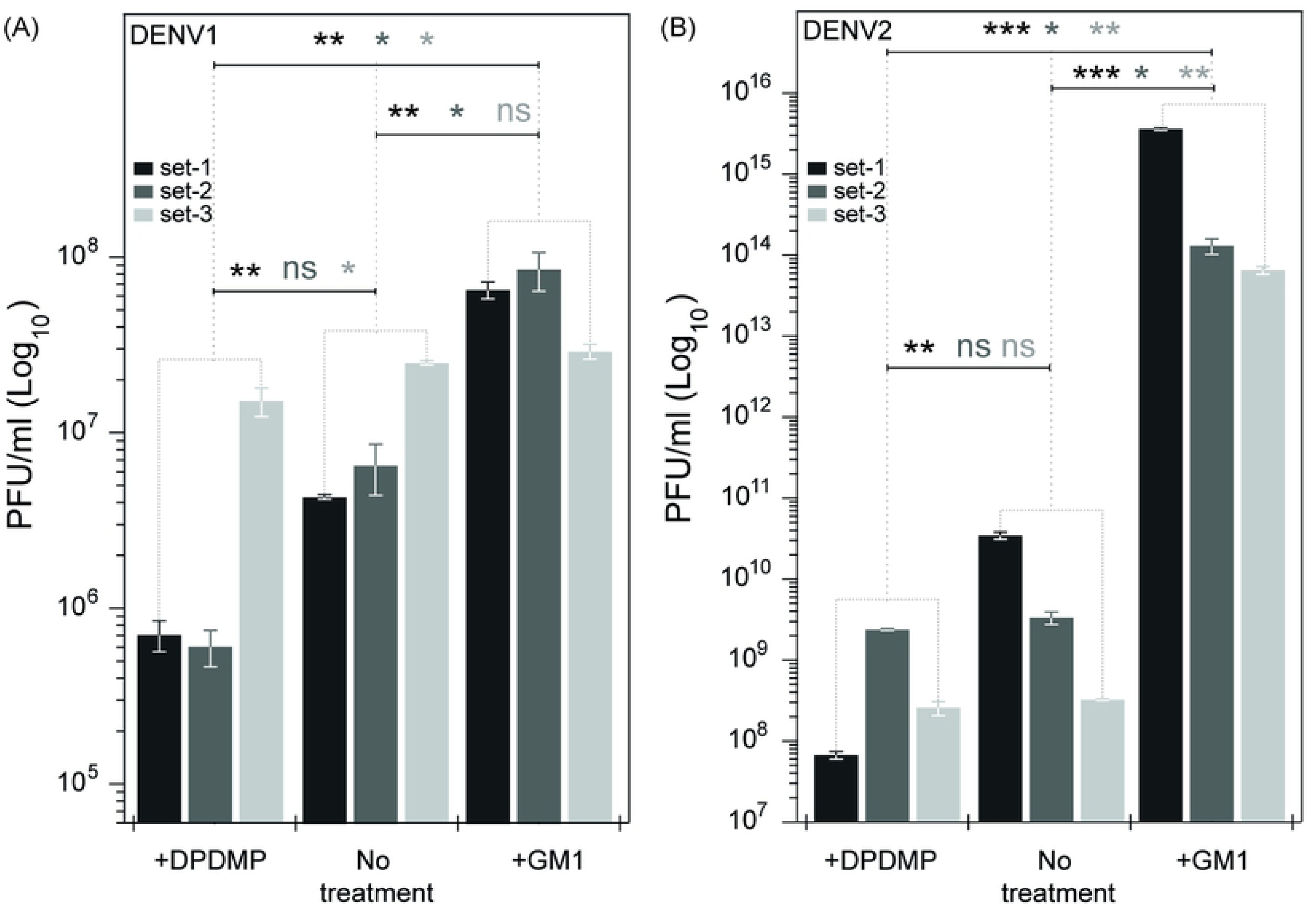
**Infectivity changes of DENV1 and DENV2 by plaque assay in presence and absence of GM1a.** (A) DENV1 with GM1a shows an increase in infectivity, while the GM1a depleted condition shows a reduction in infectivity as compared to the untreated cells. (B) DENV2 with GM1a shows a substantial increase in infectivity, while the GM1a depleted condition shows a reduction in infectivity as compared to the untreated cells. Error bars represent the SEM.

### Interaction of GM1a with DENV1 and 2 triggers increased diffusion of virus on the cell surface

The interaction of DENV with GM1a was further studied using fluorescence based SPT, where the movement of DENV1 and DENV2 was observed in GM1a depleted and GM1a enriched (cells were treated with D-PDMP and subsequently enriched with GM1a) live Vero cells. The diffusion coefficient of both DENV1 and DENV2 shows an increase in GM1a enriched as opposed GM1a depleted cells (Fig 4, S8 Fig). The Diffusion coefficients of DENV1 and DENV2 on cell membranes of GM1a depleted cells show similarity to each other at 0.005±0.001 μm^2^/s (Avg±SEM; tracks = 260) and 0.005±0.001 μm^2^/s (Avg±SEM; tracks = 487), respectively. In GM1a enriched cells, DENV1 (colocalized with GM1a) showed a diffusion coefficient of 0.010±0.002 μm^2^/s (Avg±SEM; tracks = 86), which is an increase in D from its movement in GM1a depleted cells (P = 0.0676, difference is not statistically significant at 95% confidence interval), while DENV2 showed a similar trend with an increase to 0.015±0.002 μm^2^/s (Avg±SEM; tracks = 417) in GM1a enriched cells as compared to trajectories in GM1a depleted cells (P < 0.0001, difference is statistically significant at 95% confidence interval) (Fig 4). This indicates that DENV1 and DENV2 movements are influenced by the association with GM1a.

**Fig 4.**
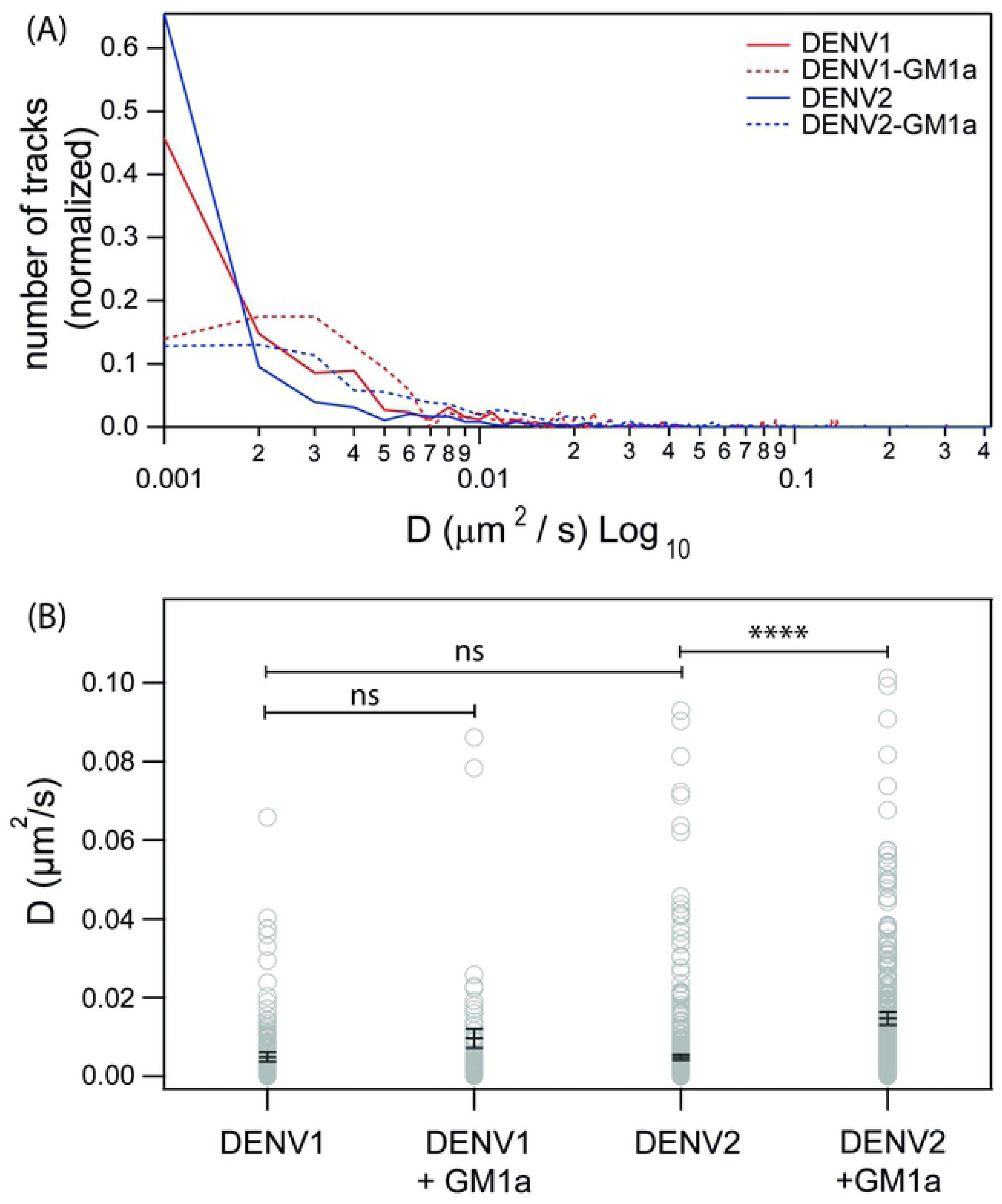
**Distribution of diffusion coefficients from MSD curves for DENV on live Vero cell membrane surface.** (A) The normalized distributions of DENV1 compared with DENV1-GM1a, and DENV2 with DENV2-GM1a. The virus once colocalized with GM1a shows a population shift toward increased diffusion coefficients (B) Average±SEM shown, with diffusion coefficient distribution of 2D SPT tracks of DENV1, and DENV2 labelled with DiI’, in presence and absence of GM1a-Bodipy. The distribution of diffusion coefficient of DENV1 increase from 0.005 ± 0.001 μm^2^/s (Avg±SEM; tracks = 258) to 0.01 ± 0.002 μm^2^/s (Avg±SEM; tracks = 86) for with and without colocalization with GM1a respectively (not statistically significant at α = 0.05), while DENV2 diffusion coefficient distribution shows a shift to from 0.005 ± 0.001 μm^2^/s (Avg±SEM; tracks = 483) to a higher D of 0.015 ± 0.002 μm^2^/s (Avg±SEM; tracks = 415) when colocalized with GM1a. Cells were pre-treated with D-PDMP to deplete GM1a for trajectories of DENV with no GM1a. Cells pre-treated with D-PDMP were enriched with GM1a-Bodipy and colocalized trajectories are presented as DENV-GM1a.

The changes in diffusion of DENV on cell membranes might also indicate a change in the mode of diffusion. GM1a is located in lipid rafts of mammalian cell membranes, and any changes in organization may influence the diffusion of cargo associated with GM1a on cell membranes [78, 79]. For this purpose, we performed ImFCS measurements on live Vero cells transfected with the raft marker GFP-GPI and pre-enriched with GM1a, to compare changes that occur before and after overlay of DENV. ImFCS is capable of determining the diffusion mode of cell membrane components [80]. The GFP-GPI *τ*_0_ values reduced from 0.53 ± 0.30 s before (cells = 6, with 3 sets of 21x21 pixel areas per cell for ImFCS) to 0.34 ± 0.15 s after DENV1 addition (P = 0.0131, difference is statistically significant at 95% confidence interval) (Fig 5B). While in the case of DENV2, a similar trend was observed where *τ*_0_ showed a reduction from 0.68 ± 0.31 s to 0.51 ± 0.30 s (cells = 5, with 3 sets of 21x21 pixel areas per cell for ImFCS) in the absence and presence of DENV2 respectively (P = 0.1382, difference not statistically significant.) (Fig 5). This reduction in *τ*_0_ for GFP-GPI raft marker indicates a change in the probe diffusion mode from transient domain confined to a freer diffusion, which in turn indicates a change of the raft organization tending towards a slightly less rigid organization allowing more freedom of movement for lipids and embedded proteins. This reduction in rigidity could be attributed to the faster movement of both DENV1 and DENV2 when colocalized with GM1a, as observed in 2D SPT trajectory data. These results were compared with ImFCS studies before and after addition of CTxB (labelled with Alexaflour555) under the same conditions. In the case of CTxB, however, there was an inversion in the GFP-GPI *τ*_0_ value, where there was an increase in *τ*_0_ from 0.58 ± 0.50 s to 1.44 ± 0.75 s (cells = 4, with 3 sets of 21x21 pixel areas per cell for ImFCS) before addition of CTxB and after the addition of CTxB (P = 0.0032, difference is statistically significant.) (Fig 5). This indicates a possible increase in the rigidity of the lipid raft after CTxB binding. CTxB is known to bind multiple GM1a receptors (up to five) and is reported to stabilize raft domains via a lipid cross-linking mechanism [81, 82], leading to an increased rigidity to the lipid raft they reside on, which could lead to the GFP-GPI probe to experience a more rigid raft environment, leading to the increase in *τ*_0_. Interestingly however, in the case of DENV1 and DENV2, the lipid raft region tends toward decreasing rigidity, indicating a difference in the way the virus binds to GM1a as compared to that of CTxB. The integrity of the plasma membrane after D-PDMP treatment and GM1a enrichment was checked by conducting FCS studies on GFP-GPI raft marker and DiIC18 liquid disordered marker, and it was observed that the diffusion coefficient of GFP-GPI was at 0.67 ± 0.49 μm^2^/s (cells/curves = 7/19) and 0.75 ± 0.32 μm^2^/s (cells/curves = 10/29) for untreated and D-PDMP treated Vero cells respectively, indicating that no changes to membrane raft regions occurred due to D-PDMP treatment (S9 Fig). Similarly, FCS of DiIC18(3) liquid disordered marker was conducted on both non-treated and D-PDMP treated Vero cells, and the diffusion coefficients were at 1.59 ± 0.77 μm^2^/s (cells/curves = 6/11) and 1.70 ± 0.63 μm^2^/s (cells/curves = 6/10) respectively. This indicates that the membrane disordered regions also do not undergo any organization changes due to the treatment with D-PDMP (S10 Fig).

**Fig 5.**
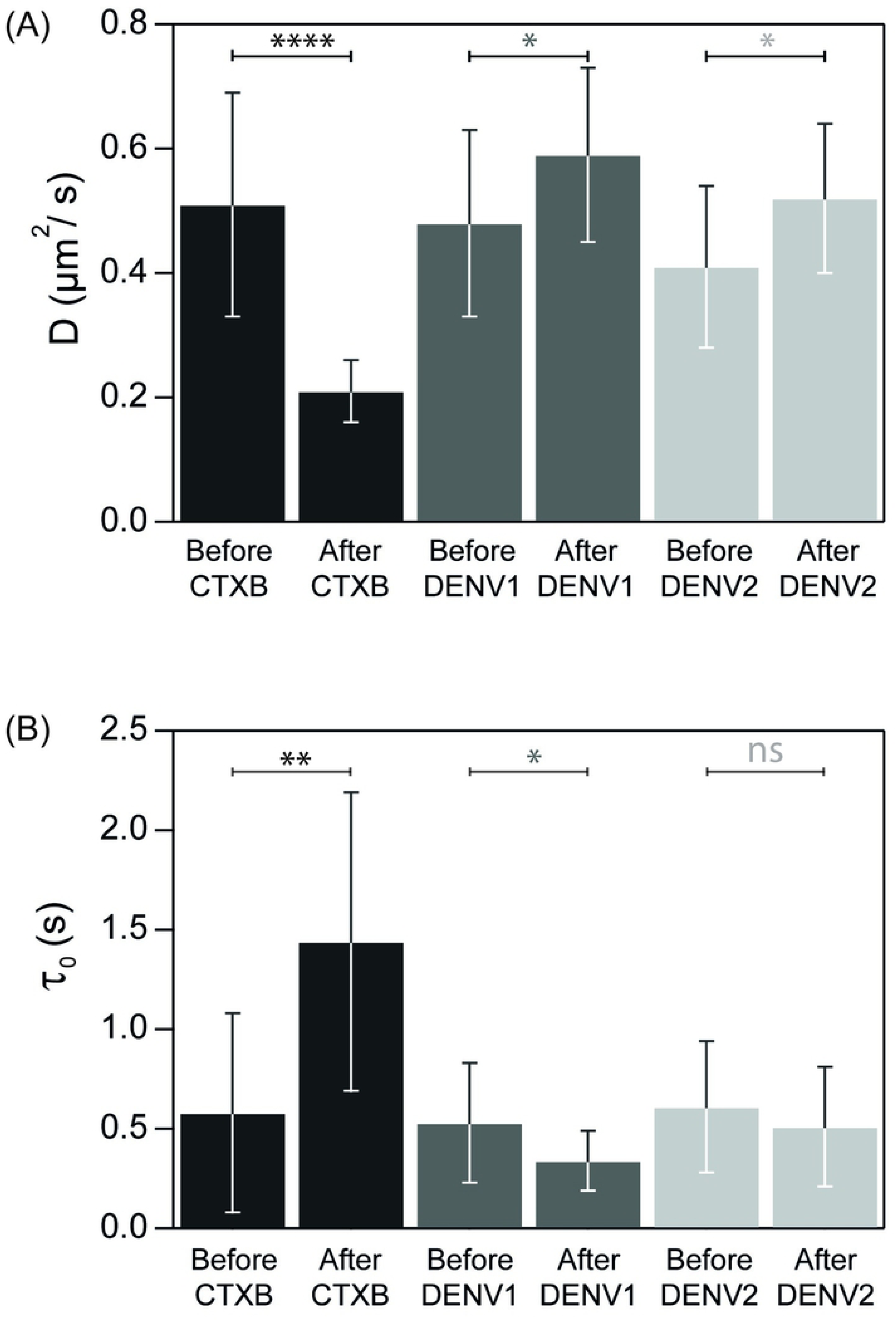
**Diffusion coefficients and Diffusion law intercepts of GFP-GPI raft probe on live Vero cell membrane measured by ImFCS.** (A) Comparison of diffusion coefficient of the raft-marker GFP-GPI, on cells treated with D-PDMP+GM1a, to observe the effect of DENV1, DENV2, and CTXB interaction with GM1a. (B) ImFCS Diffusion law intercept τ_0_ changes of the raft-marker GFP-GPI, on cells treated with D-PDMP+GM1a, to observe the effect of DENV1, DENV2, and CTXB interaction with GM1a. Error bars in both graphs represent the SD.

## Discussion

DENV is known to enter mammalian cells using many different entry mechanisms, while utilizing many different types of receptors/co-receptors/attachment factors. The range of receptors and cells DENV is reported to infect, hints towards a more ubiquitous form of entry which is available on many cells. Any virus when approaching the cell surface must find its way to these entry points for successful access into the host system. Mammalian cell membrane consists of a mosaic of lipids and proteins which are known to form liquid ordered (rafts) and liquid disordered regions, and encompass various receptors both proteinaceous and others. The cell surface is decorated with sugar antennas which form the glycocalyx, extending outward towards the extracellular environment, to regions beyond the reach of cellular receptors, which forms the first barrier that any virus will encounter as it approaches the cell surface. Thus, it is important to note that any virus that internalizes into a living cell must travel past this first barrier, and it is biologically important to identify how sugar-based molecules are involved in the virus infection process.

In this work we focus on GM1a, which is a glycosphingolipid with a sialic acid-based sugar found ubiquitously on mammalian cell surfaces, with its sugar moiety located within the glycocalyx. Sialic acid-based sugars have been widely reported to be involved in internalizing many different types of viruses, including Influenza A and SARS-Cov-2. We explore the involvement of GM1a and its sialic acid moiety in DENV internalization by using real-time fluorescence microscopy techniques. GM1a labelled with bodipy colocalizes with both DENV1 and DENV2 on live Vero cell systems, and moves along the cell surface together, indicating binding of the viral cargo onto the GM1a moiety. This binding of DENV with GM1a is further confirmed by our FCCS results, where a positive *q* value of 0.15±0.02 and 0.21±0.02 for DENV1 and DENV2 was observed. The biological significance of this binding of DENV with GM1a and its effect on infectivity of DENV1 and DENV2 was then explored by plaque assay, where it was evident that the presence of GM1a on mammalian cell surfaces significantly increases the infectivity of both DENV1 and DENV2 as compared to the GM1a depleted states. The association of DENV with GM1a shows a significant effect on the infection process of DENV, and being a ubiquitously available molecule on mammalian cell surfaces, it acts as a more universal interacting partner during virus internalization.

The sialic acid receptor GM1a is reported to bind CTxB protein cargo via the two terminal sugars (galactose and sialic acid) on the receptor, in the form of a two fingered grip, involving ionic interactions between the positively charged protein cargo and the negatively charged sialic acids, along with solvent-mediated hydrogen bonding [29]. Haemagglutinin on influenza viruses binds sialic acid receptors (commonly N-acetyl neuraminic acid (Neu5Ac)) with Avian influenza viruses preferentially binding to α2,3-linked sialic acid, and human-adapted viruses binding to α2,6-linked sialic acid moieties [44].

In this work, the binding of DENV2 virus to the sugar moiety of GM1a occurs in the regions spanning across domain I (DI) and domain II (DII) of E protein, while recombinant DENV2 E protein alone does not significantly bind GM1a under our experimental conditions. This indicates that GM1a requires quaternary contacts of a fully assembled virus to bind to the surface. The binding of GM1a sialic acid moiety may follow ionic interactions between the negatively charged sugar, and the more positively charged virus surface E protein, along with possible solvent mediated H-bonding to further stabilize the interaction. The E proteins of both DENV1 and DENV2 show sequence similarity of ∼67%, indicating preservation of similarity of E protein arrangement of the two variants. Thus, DENV1 interaction with GM1a sialic acid may show a similar binding pattern to that of DENV2.

DENV1 and DENV2 once colocalized with GM1a on live cell surfaces, shows movement along the cell membrane. Any cargo once attached to the attachment-factor/coreceptor undergoes movement along the cell surface, as it “surfs” and “tumbles” in search of the optimum orientation and supporting coreceptors and “true” receptors for its internalization. In this work, both DENV1 and DENV2 show an increase in movement speeds once it is colocalized with GM1a, as opposed to the GM1a depleted states, indicating that once attached to GM1a, the sialic acid receptor promotes faster movement of the virus on live cell surfaces. This increase in speed can allow the virus cargo to surf the cell surface more efficiently in search of its final “true” receptor. Once the true receptor is found, the virus will attain an immobile state and finally internalize. Thus, it can be interpreted that GM1a acts more as an attachment-factor or co-receptor for both DENV1 and DENV2, in assisting its internalization and infection process into live mammalian cells.

This increase in movement of DENV1 and DENV2 goes hand in hand with changes that undergo on the cell membrane organization. Once DENV associates with GM1a, there is an increase in fluidity of the membrane raft regions, indicated by the decrease in GFP-GPI raft marker *τ*_0_ value obtained by ImFCS. This indicates a change in the probe diffusion mode from transient domain confined to a freer diffusion, which in turn indicates a change of the raft organization tending towards a slightly less rigid organization allowing more freedom of movement for lipids and embedded proteins. This change in the lipid rafts possibly allow the DENV-GM1a complex to travel faster on the cell membrane surface. The interaction of DENV with GM1a, is thus different to that of CTxB binding to GM1a, as CTxB binding triggers increased cross linking between GM1a molecules inside the raft region, leading to increased rigidity. Thus, the binding of DENV with GM1a, does not appear to require cross-linking of GM1a.

GM1a may act as an important attachment-factor/coreceptor for DENV entry into mammalian cells, which support DENV movement to assist finding of the “true” receptors for virus internalization. GM1a is important in this search for the true receptor, as its absence clearly reduces the infectivity of both DENV1 and DENV2. Depletion of GM1a reduces, but does not abolish infection, and thus, GM1a is deemed to be one of many entry mechanisms into mammalian cells. The dynamics of DENV binding to GM1a need to be explored further, as the effect on the cell membrane during binding of virus to GM1a is an interesting phenomenon that is quite different to the normal way GM1a is known to participate in internalizing cargo such at CTxB. Further, the final receptor for DENV1 and DENV2 is not known as of now, and future work may enable identifying “true” receptor(s) involved in GM1a assisted entry of DENV into mammalian cells.

## Materials and Methods

### Key Resources Table

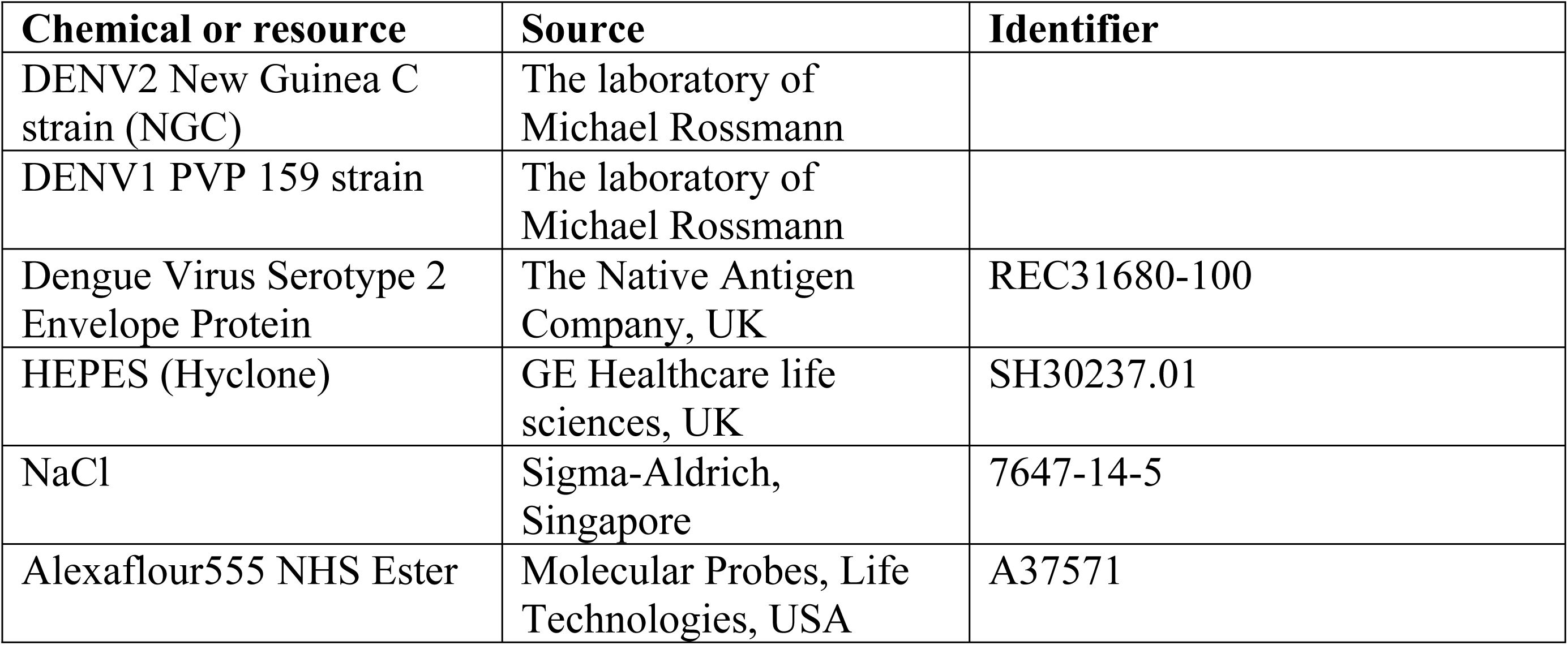
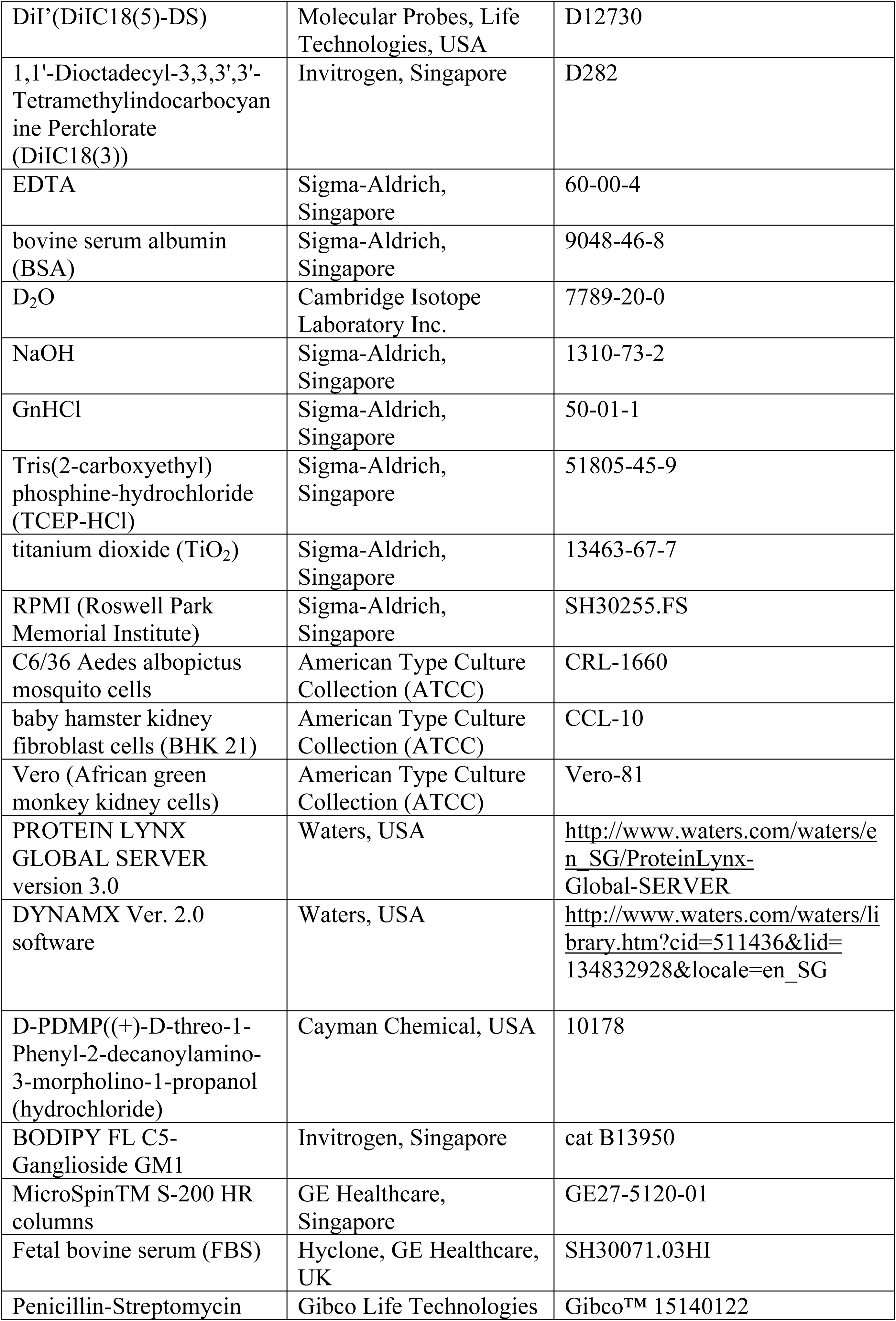
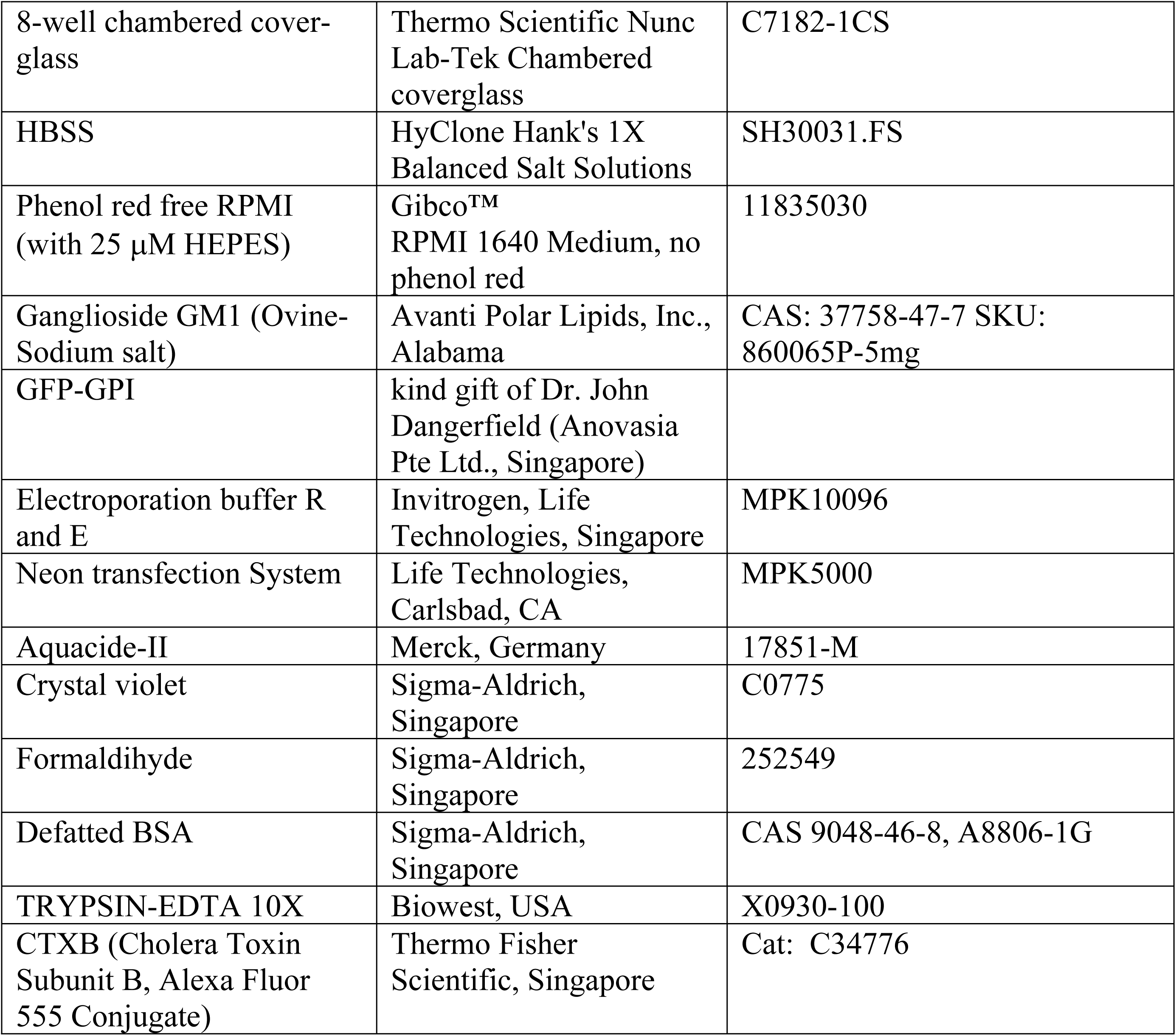

## Method Details

### Cell lines

Vero-81 (RRID:CVCL_0059), C6/36 mosquito cells and baby hamster kidney strain 21 (BHK21) fibroblast cells were from the American Type culture collection (ATCC). C6/36 were cells derived from larvae of female Aedes albopictus mosquito and were adapted to Eagle’s minimum essential medium. Cells were cloned and re-cloned by seeding single cell suspensions into petri dishes. Further information about the cell lines can be obtained through Key Resources Table.

### Cell culture

African green monkey kidney cells (Vero) and Baby hamster kidney cells (BHK21) were cultured in 10% fetal bovine serum in Roswell Park Memorial Institute (RPMI) 1640 Medium with 25 mM HEPES, and L-Glutamine, with 1% Penicillin-Streptomycin in 5% CO_2_ at 37 °C. Vero and BHK21 cells were a kind gift from Prof. Dr. Shee-Mei Lok, Duke-NUS Medical School, Singapore. Cells were passaged once they reached 90% confluence, by trypsinization (2 ml 0.25% trypsin-0.03% EDTA solution) for 2 minutes, and cells were re-seeded in new T75 flasks at a 1:5 or 1:10 split ratio with RPMI media (with 10% FBS, 1% PS, 25 mM HEPES and L-Glutamine).

### Virus preparation

DENV2 NGC and DENV1 PVP159 were produced and purified as previously described (4,83). C6/36 mosquito cells were infected with DENV2 NGC and DENV1 PVP159 at an MOI of 0.1 and grown under 5% CO_2_. The inoculant was replaced after 2 hours with RPMI medium supplemented with 2% fetal bovine serum (FBS) and allowed to incubate for 4 days, after which the virus containing supernatant was clarified from cell debris by centrifugation. The virus particles were collected by precipitation with 8% polyethylene glycol 8000 (PEG 8000) in NTE buffer (10 mM Tris-HCl [pH 8.0], 120 mM NaCl, 1 mM EDTA) and purified by centrifugation using a 30% sucrose cushion followed by a 10 to 30% potassium tartrate gradient. The virus band was collected and concentrated with a buffer exchange to NTE, using an Amicon Ultra-4 centrifugal concentrator (Millipore) with a 100-kDa molecular-mass cut-off filter to obtain glycosylated native state virus particles. The virus preparation contained low levels of contamination by immature virus as determined by test for prM with a Coomassie blue-stained SDS-PAGE. The envelope protein concentration was estimated by comparing the corresponding band with that of bovine serum albumin (BSA) at different concentrations [4,83,84].

### Viral labelling for fluorescence experiments

Purified DENV samples were labelled using either Alexa Fluor 555 NHS ester or DiI’(DiIC18(5)-DS) with ∼2.5 × 10^8^ PFU of the purified virus in 10 mM HEPES, 150 mM NaCl at pH 7.4 (HN buffer) solution. The labelling was performed at a final concentration of 750 nM and 100 nM Alexa Fluor 555 NHS ester and DiI’ (DiIC18(5)-DS) respectively (molar extinction coefficients of 71,000 M^−1^cm^−1^ and 144,000 M^−1^cm^−1^, respectively), and left at 4°C (to reduce loss of infectivity due to exposure to room temperature) for 2 hours. The Alexa Fluor 555 NHS ester labels the E protein on the virus, while the DiI’ (DiIC18(5)-DS) labels the viral bilayer [84]. The free dye molecules were filtered out by size exclusion chromatography (MicroSpinTM S-200 HR columns).

### Preparation of GM1a for cell enrichment

The GM1a-Bodipy 25 μg powder was first diluted in 50 μl chloroform: ethanol (19:1) solution, transferred to a clean 25 ml round bottom flask, dried under nitrogen gas to evaporate the solvent and further dried under vacuum for 1 hour, after which it was dissolved using 200 μl of absolute ethanol. This ethanol mixture of GM1a-Bodipy/GM1a was then mixed by vortexing into 0.34 mg/mL solution of defatted BSA in Hanks’ buffered salt solution + 10 mM HEPES, pH 7.4 (HBSS/HEPES) to make a 3 μM solution of GM1a in BSA which was aliquoted as 1 mL batches in Eppendorf tubes and stored at -20 °C, as recommended by the product information sheet. Similarly, Ganglioside GM1a solid was dissolved in a chloroform:ethanol (19:1) solution, to make a 10 mg/ml stock solution. From this a volume of 40 ml was taken and following the same protocol as above, a final stock of 5 μM GM1 in BSA was prepared and stored at -20 °C until experiment date.

### Treatment of cells with D-PDMP and enrichment of GM1a

The cells were trypsinized, and spun down in a 15 mL falcon tube at 1000 rpm for 3 minutes to form a pellet. The cells were resuspended in 5 ml RPMI (with 10% FBS, 1% PS, 25 mM HEPES and L-Glutamine) and seeded at ∼2000 cells per well in an 8-well chambered cover-glass (Thermo Scientific Nunc Lab-Tek Chambered coverglass). A solution of D-PDMP was diluted in 100% Ethanol to form a 10 mg/mL solution and diluted in 1xPBS to a final concentration of 0.5 mg/ml stock solution. This stock was diluted in cell culture media to prepare a solution of 10 μM. The cells were left to adhere to the coverglass, and after two hours, the media was replaced with media containing 10 μM D-PDMP ((+)-D-threo-1-Phenyl-2-decanoylamino-3-morpholino-1-propanol (hydrochloride)) and left for 2 days to inhibit endogenous GM1 synthesis in cells. The cells were rinsed three times with HBSS and were overlaid with 200 μl volume at a concentration of 300 nM or 100 nM GM1a-Bodipy for imaging experiments and FCS/FCCS experiments respectively in Hanks’ buffered salt solution + 10 mM HEPES, pH 7.4 (HBSS/HEPES). The GM1a-Bodipy was left on the cells for 30 minutes at 4 °C, and washed three times with phenol red free RPMI (with 25 μM HEPES). Labelled DENV was then overlaid at a multiplicity of infection (MOI) of 50 in phenol red free RPMI (with 25 μM HEPES), at 4 °C, then was washed and replaced with fresh media for imaging [85].

### Transfection protocol

For transfection with GFP-GPI, cells were trypsinized, and the volume of 1 million cells (counted by Biorad tc10 automated cell counter, Bio-Rad Laboratories, Inc Singapore) was spun down in a 15 mL falcon tube at 1000 rpm for 3 minutes to form a pellet. This pellet was resuspended in 9 μl electroporation buffer R (Neon transfection buffers), and gently mixed in with 500 ng of GFP-GPI plasmid, and transfected using the Neon transfection System with a 10 μl transfection tip using 1,200 pulse voltage (v), with 2 pulses. The cells were resuspended in 3 ml RPMI (with 10% FBS, 25 mM HEPES and L-Glutamine), and were seeded ∼2000 cells per well in 8-well chambered cover-glass (Thermo Scientific Nunc Lab-Tek Chambered Coverglass) and kept at 37 °C in 5% CO_2_.

### DiIC18(3) staining of live cell membrane

DiIC18(3) (1,1’-Dioctadecyl-3,3,3’,3’-Tetramethylindocarbocyanine Perchlorate) was dissolved in DMSO and diluted to a final DiIC18(3) concentration of 100 nM in 1 × HBSS with vortexing. Vero cells were rinsed three times with HBSS, and overlaid with 100 nM solution and left at 37 °C in 5% CO_2_ incubator for 15 minutes. The cells were then washed three times using phenol red free media and left in the same media for imaging and FCS/FCCS experiments.

### Plaque assay

BHK21 cells were grown till 90% confluency in a T75 flask, and trypsinized. The trypsinyzed cell suspension in RPMI was centrifuged to remove debris and re-suspended in 50 mL of RPMI (with 10% FBS, 1% PS, 25 mM HEPES and L-Glutamine). A volume of 1 ml of this cell suspension was seeded per well, into two 24 well cell culture plates and left for one day to allow cells to form a continuous monolayer. The virus solution was serially diluted in ten-fold dilutions, using RPMI (with 4% FBS, 25 mM HEPES and L-Glutamine). The media was aspirated off of the cell monolayer, and a volume of 100 μl of each dilution was added in triplicate to the wells. This was left to incubate at 37 °C with 5% CO_2_, for 2 hours with plate tilting every 15 minutes to stop drying of cells. The virus overlay was then aspirated out, and a viscous overlay made up of RPMI with 1% Aquacide-II and 2% FBS was placed over the cell monolayer, and left to incubate for 7 days. The plates were tipped to remove the viscous overlay, and stained using few drops of crystal violet (0.5% (wt/vol) crystal violet–25% formaldehyde), and was left for 1 hour, before washing under a running tap to remove unstained plaque areas. The plaques were counted by visual inspection and the PFU/ml was given by the (average number of plaques/ (Dilution factor ×Volume of virus added (mL))).

### Confocal microscope setup

An FV1200 confocal microscope (Olympus, Tokyo, Japan) equipped with a time-resolved FCS upgrade kit (PicoQuant, Berlin, Germany) was utilized in this work. The confocal was equipped with a pulsed 485 nm laser (LDH-D-C-488, PicoQuant, Germany) operated at 20 MHz repetition rate, a 543 nm continuous wave (cw) laser (GLG 7000, Showa Optronics, Japan), 488 nm cw laser, and 635 nm cw laser. All lasers were operated at 5 μW power before the objective, on live cell membranes and passed through a 60x, NA 1.2 water immersion objective (UplanSApo, Olympus, Japan), while the fluorescence emission is routed through a 405/488/543/635 dichroic mirror (Chroma Technology), confocal pinhole of one airy unit, and band-pass emission filters 513/17 (Brightline; Semrock, IDEX Health & Science, LLC, New York) and a 615/45 (XF3025 32833, Omega Optical, USA) for green and red emissions in FCS and FCCS experiments. Band pass filter of BA505-525, and BA655-755, for green and red respectively was utilized for imaging and SPT experiments. The emission signal was detected by an avalanche photodiode (SPCM-AQR14; PerkinElmer). The photon counts from the detector were registered by a TimeHarp 260 time-correlated single photon counting board (PicoQuant) and processed by the SymPhoTime 64 software (PicoQuant) [86, 87].

### FCS measurements on confocal microscope

FCS was performed on Vero cells at 37 °C, in phenol red free media with 25 μM HEPES on a FV1200 confocal microscope (Olympus, Tokyo, Japan). FCS measurements in the green channel for GM1a-bodipy and GFP-GPI was performed by illuminating the sample with a pulsed 485 nm laser. FCS in the red channel for DiIC18(3) labelled cell membranes were performed by illumination with a 543 nm continuous wave (cw) laser. The signal was collected at detector without a beam splitter using a single detection channel and correlation functions were calculated using SymPhoTime 64 software (PicoQuant). Determination of dimensions of the effective detection volumes were performed by calibration of FCS measurements in solutions of reference dyes atto488 and atto565 (Atto-Tec, Siegen, Germany) of green and red channels respectively. The diffusion coefficient of Atto dyes was taken as 400 μm^2^/s at room temperature, based on previously reported values [86–88].

### Fit models for confocal FCS measurements

For FCS in the confocal, in the simplest case, fluorescent molecules moving in 3D diffusion in the confocal volume is fitted using a single particle (1p) 3D diffusion fit model.

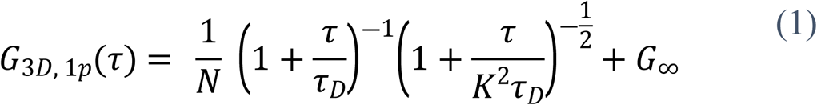

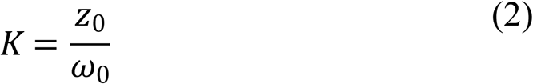

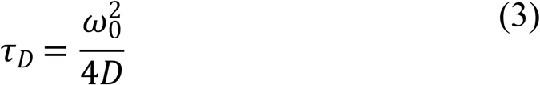

where *K* is the structure factor which defines the shape of the observation volume and *τ*_*D*_ is the diffusion time.

In the case of 2D diffusion on cell membranes, the fitting model can be simplified as

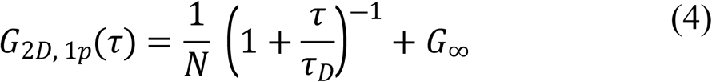

If there are multiple diffusion components in the system, the linear sum of all these individual components weighted with their respective mole fractions, gives the *G*_*τ*_, and can be expressed as

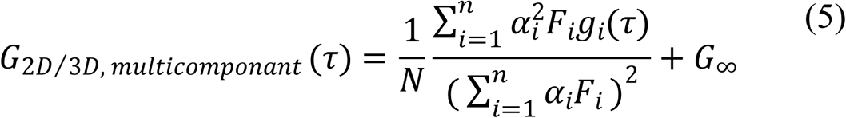

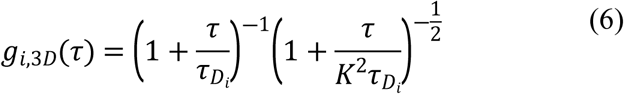

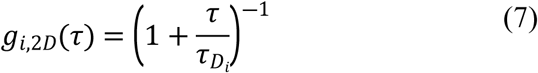

Where *α*_*i*_ is the ratio of the brightness of the *i*^*th*^ species to that of species 1. *τ*_*D*_ is the diffusion time and *F*_*i*_ is the mole fraction of the *i*^*th*^species.

The overall ACF is given as the product of all the individual dynamic processes that are present in the system, including the triplet state relaxation of the fluorophore, and can be written as

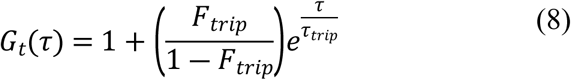

where *F*_*trip*_ is the fraction of the triplet state and *τ* _*trip*_ is the relaxation time of the triplet state. The fitting models for 2D and 3D diffusion with triplet contribution is as below

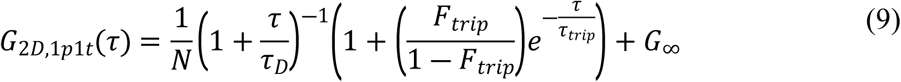

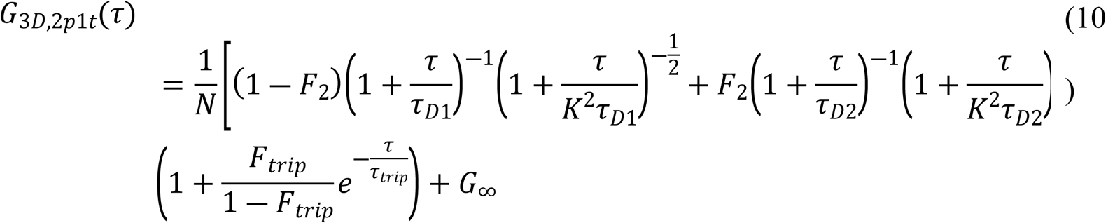

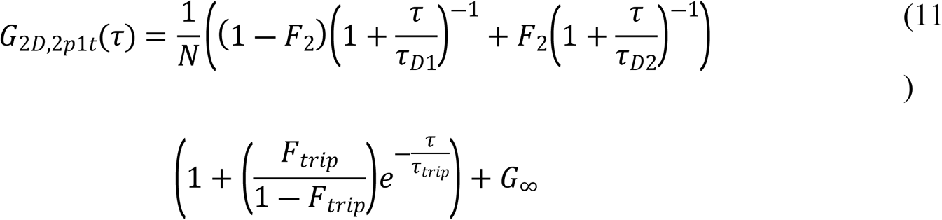

### FCCS on Confocal microscope

FCCS was performed on Vero cells at 37 °C, in phenol red free media with 25 μM HEPES on a FV1200 confocal microscope (Olympus, Tokyo, Japan). Quasi-PIE FCCS was performed by using pulsed 485 nm and the 543 cw laser to illuminate sample simultaneously. The emission was split by a 560DCXR (Chroma Technology, Bellows Falls, VT) emission dichroic filter cube to allow green and red emission to travel to the two separate detectors. The after pulsing and spectral cross talk was corrected by statistical filtering, using the FLCS script for spectral crosstalk removal via FLCCS on SymPhoTime 64 software (PicoQuant) [86, 87].

### Confocal FCCS *q* value evaluation

The degree of cross-correlation, *q* (the ratio of the concentration of the double-labelled species to that of all the particles carrying the less abundant label), was calculated as in equation (12), where the increase in *q* indicates a higher level of interaction between the two biomolecules [88].

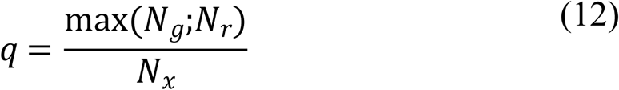

The derivation of *q* factor is as below;

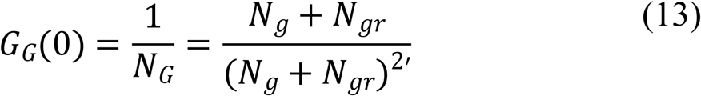

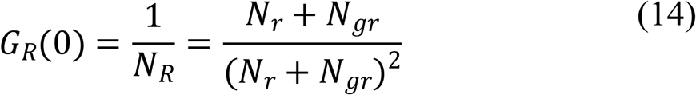

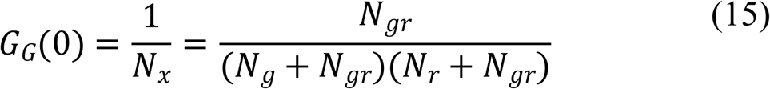

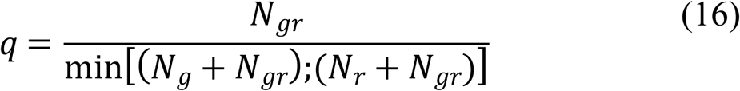

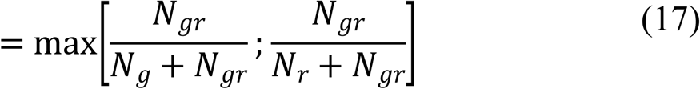

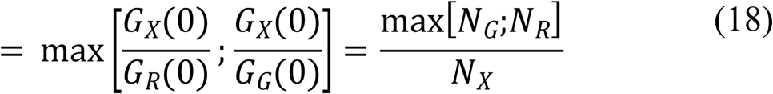

*N*_*g*_, *N*_*r*_ and *N*_*x*_ are the background corrected particle numbers extracted from the ACFs in the green and red channel and from the CCF, respectively. Ten cell measurements with three 100 s acquisitions each where measured and the standard deviations of the *q* mean was calculated for all measurements. The number of individual cells and measurement statistics are provided in the results section.

### 2D SPT of DENV on live cell membrane and imaging on Confocal

Vero cells were transfected with GFP-GPI and seeded in 8-well chambered cover-glasses. DENV labelled with DiI’ (DiIC18(5)-DS) and overlaid on cells at 4 °C for 10 minutes. 2D SPT was performed by acquiring time lapse image series of labelled DENV on Vero cells at 37 °C, in phenol red free media with 25 μM HEPES on a FV1200 confocal microscope (Olympus, Tokyo, Japan). Sample was illuminated with 488 nm laser for green probes, and excited with 635 nm laser in the red. The emission was passed through band pass filter of BA505-525, and BA655-755, for green and red respectively before being captured by the PMT. 512x512 pixel frame size at a sampling rate of 2.0 μs/pixel (2.26 s/frame), and at a zoom level of x3, with *xy* resolution of 0.137 μm/pixel, and *z* resolution of 1 μm, for 75 frames. The time-lapse image series was analysed using ImageJ plug-in mosaic [89] (MOSAIC Group, Centre for Systems Biology Dresden (CSBD), Dresden, Germany), to do particle localization and trajectory linking. The trajectories were extracted and analysed using the MATLAB class @msdanalyzer (built and maintained by Jean-Yves Tinevez, Institut Pasteur, France) [90] to generate MSD curves of trajectories, after correction for drift using velocity autocorrelation. These MSD curves were fitted using Igor Pro (WaveMetrics, Portland, OR) to obtain the diffusion coefficients of trajectories, in the case of Brownian motion, and velocities were obtained in the case of active transport.

### ImFCS (Imaging FCS) setup on TIRF microscope

ImFCS was performed on an Olympus Inverted epi-fluorescence microscope 1X83 equipped with a motorized TIRF illumination combiner (cell^TIRF/IX3-MITICO, Olympus) which allows simultaneous illumination with four laser lines. The system uses a UApoN 100x/1.49 Olympus oil-immersion objective, the excitation laser was passed onto a ZT 405/488/561/640rpc (Chroma Technology, USA) dichroic mirror, to reflect the laser light on the back focal plane of the microscope objective. The laser light incident angle was adjusted to TIRF mode by help of the Olympus Xcellence software. The GFP-GPI in cell sample was excited with the 488 nm excitation laser (Olympus Cell laser) at 100 μW before the objective, and sample signal was acquired by collecting the fluorescence emission (after passing through the ZT 405/488/561/640rpc dichroic mirror), by filtering through a laser quad band ZET405/488/561/647m (Chroma Technology, USA). The signal was recorded on an EMCCD Andor iXon3 X-9388 EMCCD camera (128x128 pixels, 24 μm pixel size), for 50000 frames at 0.002 s exposure time per frame. The microscope was fitted with a CO_2_/Air gas chamber (Live Cell Instrument, FC-5, Chamlide, Seoul, Korea) and sample stage was maintained at 37 °C, with 5% CO_2_ for all cell measurements.

### ImFCS on live cell membranes

Vero cells where transfected with GFP-GPI and treated with D-PDMP for two days to deplete GM1. The cells were then enriched with GM1a. ImFCS measurements were carried out and diffusion law intercept was determined on the same cell for before and after addition of DENV1, DENV2 and CTXB. DENV1, DENV2 and CTXB were labelled with Alexa fluor 555, and the cells measured after addition of DENV1, DENV2 or CTXB were first checked with 561 nm laser illumination (Olympus Cell laser), to confirm the presence of DENV1, DENV2 or CTXB on the cell of interest. A region of interest (ROI) of 21 × 21 pixels (5 × 5μm^2^) was selected from acquired image stacks away from cell edges for ImFCS and diffusion law analysis. The ACFs for all pixels in the image were calculated using a multi-tau correlation scheme [80], and the signal was corrected before fitting, with an exponential of polynomial bleach correction [91]. The ACFs were fitted using the model in equation (19) on the home written ImFCS plugin which runs on ImageJ, to generate diffusion coefficient (D) maps and diffusion law plots.

### Imaging FCS diffusion law

FCS diffusion law [92, 93] exploits the space dependent property of membrane diffusion to explore and identify the mode of membrane organization in living cells. It is capable of differentiating between free diffusion, diffusion in domain partitioning and meshwork environments. Imaging FCS (ImFCS) [80,94,95] is the adaptation of FCS diffusion law, where the dependence of diffusion time of the fluorophore (τD) on observation area (*A*_*eff*_) is described by *τ*_*D*_(*A*_*eff*_) = *τ*_0_ + ^*A*^*eff* ^*D*^. *A*_*eff*_ is given by the convolution of the detection area (or pixel area) with the point spread function and different *A*_*eff*_ values were obtained by binning the pixels post-acquisition (n × n with n = 1–5). The magnitude and sign of intercept *τ*_0_ indicates the mode of membrane diffusion and state of membrane packing. In a diffusion law plot, free diffusion produces a straight line that passes through the origin with *τ*_0_ values of 0.0 ± 0.1 s [96]. Heterogeneous systems with domains or meshwork producing different diffusion law plots with characteristic positive and negative *τ*_0_ values for domain partitioning and meshwork diffusion, respectively [94, 97].

### Fitting of FCS curves in ImFCS

FCS curves in ImFCS was fitted using equation;

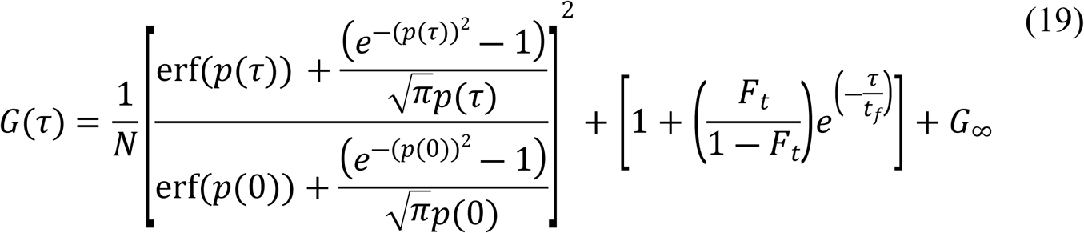

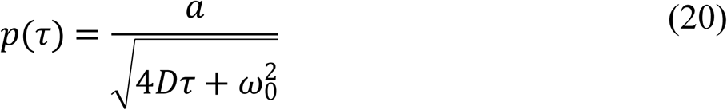

Where *G*(*τ*) is the temporal autocorrelation function, *N* is the number of particles, *a* is the pixel size and *ω*_0_ is the 1/*e*^2^ radius of Gaussian approximation of the microscope point spread function. *F*_*t*_ is the fraction of particles in triplet state, *t*_*f*_ is the average time molecules spend in the triplet state, *G*_∞_ is the convergence of *G*(*τ*) at long lag times. The fitting parameters were *N*, *D*, *F*_*t*_, *t*_*f*_ and *G*_∞_.

### DENV overlay on live Vero cells for FCS, FCCS and ImFCS

Vero cells were seeded and treated with D-PDMP to deplete cells of endogenous unlabelled GM1(99), and enriched with 50 nM GM1a-Bodipy, and overlaid with an MOI of 500 DENV1 or DENV2 for FCCS experiments. The DENV was labelled with Alexa fluor 555 NHS. FCCS was conducted at 37 °C in phenol red free HEPES supplemented imaging media. The ACFs and the CCFs where calculated after FLCCS correction for detector cross-talk, and *q* values were calculated.

### DENV overlay on live Vero cells for imaging and 2D SPT

Vero cells were plated on 8-well chambered cover-glass (Thermo Scientific Nunc Lab-Tek Chambered coverglass) after necessary dilution in media, and treated for different experimental conditions (D-PDMP/transfection/DiI staining/GM1a-Bodipy enrichment) as required. DENV was labelled and filtered, and diluted in phenol red free media with 25 μM HEPES to obtain an MOI of 50, and overlaid on cells which were washed three times using phenol red free media. This was left at 4 °C for ten minutes to help virus reach the live cell membrane and to halt cellular endocytosis [99]. The 8-well chambered cover slides were then transferred to the microscope stage for imaging.

### Deuterium labelling and quench conditions for Amide hydrogen deuterium exchange mass spectrometry

Purified dengue virus solubilized in NTE buffer at pH 8.0 was incubated at 37°C in PBS buffer reconstituted in D_2_O (99.90%) resulting in a final D_2_O concentration of 90%. For generating DENV2: GM1a complex, viral particles were preincubated at a molar ratio of 1: 125 (E protein monomer: GM1a sugar) at 37°C for 30 min prior to each deuterium exchange reaction. Deuterium labelling was performed for 1 minute for both free and GM1a bound state followed by quenching the exchange reaction by adding prechilled quench buffer. Quench buffer contained 1.5 M GnHCl and 0.25 M Tris(2-carboxyethyl) phosphine-hydrochloride (TCEP-HCl) and after adding quench buffer the solution was incubated at 4 °C on ice for 30 sec followed by addition of titanium dioxide to precipitate envelope lipids. Precipitated envelope lipids were removed before injecting the sample for pepsin digestion using 0.22 µm centrifugal filters at 10000 rpm for 1 min. Deuterium exchange was also performed on free soluble E protein (1mg/ml, C-terminal (His)_6_ tag) and sE protein: GM1a (1:50) for labelling times of 1, 10- and 100-min. sE protein was reported to exist dominantly in a monomeric and dimeric state at 37°C and 22°C respectively [101]. HDXMS was carried out at 22°C to map GM1a interactions to the dimeric state of E protein. Same quench conditions were used for both sE protein states with the exception that E protein from intact Dengue was treated with TiO_2._This was not required for recombinant sE protein as it lacks stem helices and lipid membrane.

FCCS experiments have shown colocalization of GM1a glycosphingolipid with DENV2 and DENV1 showing the stable binding of GM1a to the viral surface. In the absence of equilibrium dissociation constant (K_d_), we used high molar ratio of GM1a sugar moiety to E protein (indicated above) in our deuterium exchange reactions. We observed no significant changes in sE protein (monomer) in the presence of GM1a at 1: 50 molar ratio either due to weak binding (high K_d_) or lack of quaternary contacts akin to a virion. Also, no bimodal mass spectra was observed in sE protein: GM1a state to suggest partial or unsaturated binding of GM1a at virus E protein. Hence, in subsequent deuterium exchange reaction with DENV2 we used a higher concentration of DENV2 (E protein monomer) to GM1a molar ratio (1:125). Peptides showing protection in the presence of GM1a (DENV2: GM1a state) show unimodal distribution of mass spectra which suggests uniform binding across viral particle.

### Mass Spectrometry and peptide identification

Quenched samples of 35 pmol and 100 pmol from DENV2 and sE protein (free and GM1a complex states) were injected onto nanoUPLC HDX sample manager (Waters, Milford, MA) respectively. Injected samples were proteolyzed in online mode using pepsin immobilised Waters Enzymate column (2.1 × 30 mm) in 0.1% formic acid in water at a flow rate of 100 μl/min. The proteolyzed peptides were trapped in a 2.1 × 5 mm C18 trap (ACQUITY BEH C18 VanGuard Pre-column, 1.7 μm, Waters, Milford, MA). Elution of pepsin digested peptides was performed using acetonitrile (ACN) gradient of 8 to 40 % in 0.1 % formic acid at a flow rate of 40 µl min^-1^ into reverse phase column (ACQUITY UPLC BEH C18 Column, 1.0 × 100 mm, 1.7 μm, Waters) pumped by nanoACQUITY Binary Solvent Manager (Waters, Milford, MA). Peptides were ionized using electrospray ionization mode and sprayed onto SYNAPT G2-Si mass spectrometer (Waters, Milford, MA). HDMS^E^ mode acquisition and measurement was employed. 200 fmol μl^−1^ of [Glu^1^]-fibrinopeptide B ([Glu^1^]-Fib) was injected at a flow rate of 5µl/min into mass spectrometer for calibration and lockspray correction.

Protein Lynx Global Server v3.0 (PLGS v3.0) was used to identify the peptides from undeuterated mass spectra (HDMS^E^). Search for peptide identification was performed on sequence database of dengue 2 NGC strain with E, M and C protein. No specific protease and variable N-linked glycosylation modifications were chosen in search parameters to carry out the sequence identification. Peptide identification parameters intensity of 2500 for product and precursor ions, minimum products per amino acids of 0.2 and a precursor ion mass tolerance of <10 ppm using DynamX v.3.0 (Waters, Milford, MA). Peptides present in at least two out of three undeuterated samples were retained. Reported deuterium exchange values are uncorrected for back exchange and all the reactions are performed in triplicates. Three technical replicates were carried out for each deuterium exchange reaction for all states and their average values were used to generate the plots. Standard deviation within ±0.5 Da was observed for all the peptides. Therefore, a deuterium exchange difference of ±0.5 Da was chosen as significance threshold. List of peptides identified in the current HDXMS experiments is shown Supplementary file 1.

## AUTHOR CONTRIBUTIONS

S.N.T. and T.W designed, analyzed, and interpreted the results. S.N.T performed and interpreted the FCCS, FCS, SPT, ImFCS, and biological assays. P.V.R. performed and interpreted the HDXMS experiments. J.C.W.B. produced and supplied virus samples. S.N.T., P.V.R., G.A. and T.W wrote the manuscript.

## Supplementary Figure legends

**S1 Fig. Controls for FCCS experiments**.

FCCS control experiment to confirm no unfiltered dye interferes with the FCCS results. FCCS curves for Spin filtered solution resulting from starting solution of 500 nM concentration Alexa flour 555 on cells incorporated with GM1a-Bodipy gives no signal in the red channel and no cross correlation, indicating no excess dye is present under the conditions used for labelling DENV. Cells were pre-treated with DPDMP before enriching with GM1a

**S2 Fig. Colocalization of GM1a with DENV1, DENV2 and CTXB on VERO cell surface.**

(A) Gm1a-Bodipy colocalized with DENV1-DiI’. (B) Gm1a-Bodipy colocalized with DENV2-DiI’. (C) Gm1a-Bodipy colocalized with CTXB-555. (D) The control with Red and Green Beads shows no colocalization. Scale bar: 5 mm. Statistical data is given in S1 Table.

**S1 Table: Colocalization analysis of virus/CTXB and GM1a in live mammalian cells.** Mander’s coefficients of colocalization and Costes P-values were determined for DENV1, DENV2 and CTXB colocalization with GM1a.

**S3 Fig. Infectivity changes of DENV1 and DENV2 by plaque assay on BHK21 cells in presence and absence of GM1a.** DF = Dilution factor

**S4 Fig. Difference Plot.**

Differences in average number of deuterons exchanged (Y-axis) in DENV2 E protein:GM1a state and free DENV2 E protein state is plotted in a deuterium uptake difference plot for t = 1, 10 and 100 min labelling times. Pepsin proteolysed peptides are listed from N to C terminus (X-axis). DENV2 E protein shows no protection in the presence of GM1a and shows increases in exchange in the peptide regions as labelled. Standard deviations are shaded gray and plots were generated using DynamX 3.0 software. The data is uncorrected for maximum deuterium content of 90% under experimental conditions, and unadjusted to accommodate loss in deuterons exchanged due to back-exchange (average back-exchange ∼15% under our HDXMS conditions).

**S5 Fig. Sequence alignment of DENV1 and DENV2 E protein.**

**S6 Fig. Coverage maps.**

(A) Coverage map of E protein from dengue 2 NGC. 36 peptides spanning 63% of E protein sequence. (B) Coverage map of E protein from soluble dengue 2 NGC E protein. 107 peptides spanning 80% of E protein sequence.

**S7 Fig. Infectivity of DENV1 and DENV2 by plaque assay for virus labelling conditions.**

(A) Plaque assay wells for DENV1. (B) Plaque assay wells for DENV2. (C) Infectivity levels comparison for (i) DENV1 and (ii) DENV2 (avg±SEM). P values at α = 0.05

**S8 Fig. 2D-SPT trajectories of DENV movement on live Vero cell surface.**

(A) DENV1 in the absence of GM1a (i) trajectory on live cells, (ii) MSD curve. (B) DENV1 in the presence of GM1a (i) trajectory on live cells, (ii) MSD curve. (C) DENV2 in the absence of GM1a (i) trajectory on live cells, (ii) MSD curve. (D) DENV2 in the presence of GM1a (i) trajectory on live cells, (ii) MSD curve. (Scale bar = 2.5 μm).

**S9 Fig. Diffusion coefficients of raft-marker GFP-GPI probe on live Vero cell membrane measured by FCS.**

(A) Diffusion coefficient of GFP-GPI raft probe on non-treated (19/7 = curves/cells), D-PDMP treated for GM1a depletion (29/10 = curves/cells) compared to when enriched with GM1a (25/10 = curves/cells). (B) Respective FCS curves for (i) non-treated, (ii) D-PDMP treated cells and (iii) D-PDMP and GM1a treated cells (avg±SD). Red asterisk shows the average value of the distribution.

**S10 Fig. Diffusion coefficients of DiIC18(3) non-raft probe on live VERO cell membrane measured by FCS.**

(A) Diffusion coefficient of DiIC18 on non-treated (11/6 = curves/cells) versus D-PDMP treated for GM1a depletion (10/6 = curves/cells). (avg±SD) (B) Representative FCS curves for (i) non-treated and (ii) D-PDMP treated cells. Red asterisk shows the average value of the distribution.

**S11 Fig. Properties of 2D-SPT trajectories.**

Histogram of trajectory length, with information on trajectory average diffusion coefficient (Avg±SEM) and number of trajectories included in study for (A) DENV1 in the absence of GM1a, (B) DENV1 in the presence of GM1a, (C) DENV2 in the absence of GM1a, (D) DENV2 in the presence of GM1a.

